# Integrative Responses of Leaf-Cutting Ants to Temperature Rises

**DOI:** 10.1101/2020.08.04.236844

**Authors:** Cleverson de Sousa Lima, André Frazão Helene, Agustín Camacho

## Abstract

Thermal variation has complex effects on organisms and they deal with it by combining behavioral and physiological thermal tolerance. However, we still do not understand well how these two types of traits relate to body condition (e.g. size, hydration) and environmental variables (e.g. relative humidity), some of which are typical aspects of thermal tolerance experiments (warming rates, start temperature). We explored these interactions using a set of experiments that sequentially measure behavioral (Voluntary Thermal Maxima) and physiological thermal tolerance (Critical Thermal Maxima) for individuals of *Atta sexdens rubropilosa* (Forel, 1908). We found non-linear effects of body size on behavioral thermal tolerance and refuted the traditional hypothesis that body size increases ant’s physiological thermal tolerance. Hydration state and humidity had complex effects on behavioral and physiological tolerance. However, both tolerance measures increased with heating rates and start temperature. Our work helps understanding how an ectotherm integrates stimuli affecting its thermal tolerance to decide which temperatures to avoid. We discuss implications for the ecology of ants, their labor division, and for their susceptibility to climate warming and drought.

**Summary Statement:** Here we show how internal (body size, hydration level) and external factors (heating rate, relative humidity) affect leaf-cutting ants behavioral and physiological responses to temperature rises.

## 1 Introduction

The climate crisis (Pachauri et al., 2014) makes it necessary to understand how organisms will respond to thermal stress in different contexts. While many studies have regarded variation in different parameters of organisms’ thermal tolerance (e.g. Angilletta et al., 2007; Cerdá et al., 2002; Christian & Morton, 1992), a fundamental and less understood aspect of this problem concerns how organisms integrate behavioral and physiological thermal tolerance to cope with temperature rises at the microhabitat scale.

Rises in environmental temperature have non-linear effects on physiological functions of ectothermic organisms (Angilletta, 2009; Camacho et al., 2018; Huey and Stevenson, 1979), and neural changes precede heat coma and CTmax (Jørgensen et al., 2020). At tolerable thermal levels, different aspects of physiological performance can be optimized within specific intervals (stamina, sprint speed, Huey, 1984). Nonetheless, excessively high temperatures will block these physiological processes and can kill animals in a short time (Angilletta et al., 2007; Christian and Morton, 1992; Ribeiro et al., 2012), for example, when animals attain their Critical Thermal Maxima (CTmax, Cowles and Bogert, 1944). At temperatures close to such levels, animals often move away from heat sources, exhibiting a Voluntary Thermal Maximum (VTM, Cowles & Bogert, 1944; Camacho et al., 2018) that can be conserved within species across their geography and history (Camacho & Rusch 2017, Wiens et al 2019).

Assuming that the VTM is, as CTmax is, a physiological thermal tolerance related effect, it is reasonable to expect that internal and environmental conditions of an organism must define the behavioral thermal avoidance. For example, since thermal tolerance of ants (i.e. their CTmax) can be affected by several factors (e.g. Ribeiro et al 2012), it could be expected that behavioral thermal tolerance dynamically responds to them, assuming that behavior has been selected to maintain organisms within optimal, or even lower thermal levels (Huey & Martin, 2009). Factors that may alter physiological thermal tolerance can be divided into traits of the organism and of the environment. For instance, body size is an overarching trait that affects many physiological (Jensen and Nielsen, 1975; Hurlbert et al., 2008) and ecological traits (Kaspari, 1993; Johnson, 2008; Baudier et al., 2015). This trait is known for raising the critical thermal limits of vertebrate and invertebrate ectothermic species (Angilletta et al., 2007; Christian and Morton, 1992, Ribeiro et al., 2012). Larger bodies have lower water loss rates, can store more water, and, in small arthropods’ case (i.e. ants), may help separating the body from heated surfaces, if associated to longer limbs (Kaspari and Weiser, 2002). Nonetheless, few data exist regarding the relationship between CTmax and body size in insects (e.g. Ribeiro et al., 2012; Verble-Pearson et al., 2015). In *Atta sexdens*, body size present large differences among different casts, which execute different tasks within a colony (Hölldobler and Wilson, 1990). In this way, observing body size effects on thermoregulation and thermal tolerance might help to understand constrains on the engagement on different tasks.

The level of body hydration (HL) can be another organismal trait regulating thermal tolerance among ectothermic animals. Body water loss rates increase with rising body temperatures (Edney, 1977; Lighton and Bartholomew, 1988) and lower the CTmax and VTM of some ectotherms (e.g. Anurans, Anderson and Andrade, 2017). Small arthropods also react to water stress (Denlinger and Yocum, 1998), by foraging closer to their refuges (Lighton et al., 1994) or selecting resources with higher water content (Bowers and Porter, 1981), or recruiting individuals with specific smaller size to transport water to the anthill (Ribeiro and Navas, 2008). Knowing whether dehydrated individuals lower their VTMs to protect themselves from lower CTmax values should help us understand how dehydration may trigger thermally induced changes in microhabitat use and activity (Rozen□Rechels et al, 2019). Yet, to our knowledge, such effect of dehydration has not been detected for arthropods.

Apart from these traits, some environmental conditions may also increase the risks of overheating and dehydration, and, consequently, induce changes in either behavioral and physiological thermal tolerance. For example, higher heating rates could lead an organism (ex. an ant) to seek thermal refuge at lower temperatures, in anticipation of a higher risk of exceeding their CTmax. In turn, a slower heating rate, involving longer exposure to sublethal temperatures, might also induce greater physiological stress, especially if that happens within a more drying environment. If such conditions lower an organism’s CTmax, is possible to expect a decrease in voluntary thermal maxima, under the above presented premise of selection on behavior. These conditions can be manipulated by controlling heating rates (HR) and start temperatures (ST), and the air relative humidity (RH), during thermal tolerance assays (Terblanche et al., 2007; Camacho et al., 2018). Thus, observing their effect on serial measures of the VTM and CTmax should help us better appreciate the integration of behavioral and physiological thermal tolerance, and how our experimental set ups influence such understanding.

The small size and big number of ants make them useful experimental models for thermal tolerance studies. Leaf-cutting ants *Atta sexdens rubropilosa* (Forel, 1908) also present the particularity that their casts differ in size and tasks. Thus, observing thermal tolerance changes with body size and environmental conditions, might shed light on their labor division. Herein, we tested if sequentially measured VTM and CTmax of leaf-cutting ants, were influenced by (1) body size, (2) the start temperature and heating rate, and (3) a combination of body hydration level and air’s relative humidity.

## 2 Materials and Methods

### 2.1 Study animals

Ants were supplied by the Laboratório de Ciências da Cognição (Department of Physiology, Institute of Biosciences, University of São Paulo). Six adult colonies were used. Specifically, our ants come from six colonies, raised in the laboratory at 24 °C and 75% ~ 85% relative humidity, and fed every day with leaves of *Acalypha spp*.

### 2.2. The Thermal Tolerance Meter

We developed a device capable of sequentially measuring ants’ VTM and CTmax in about 15 minutes (Fig. 1). In this device, four animals are simultaneously heated in individual identical chambers with a small thermal refuge, inserted within a thermal bath. Ants’ body temperature is monitored by a T-type thermocouple (1mm diameter, Omega ©), inserted in a freshly dead ant. That ant is always similarly sized to the ones being heated and its thermocouple is connected to a computer through a datalogger (Picolog^®^ TC H8). After the onset of warming, each individual’s VTM is registered as the body temperature measured when the ant enters and remain in the thermal refuge for at least 7 seconds. As temperature keeps raising within the thermal refuge, ants move back to the heating tube and enter thermal equilibria with the other ants and the model (see supplementary material file S1). Heating continues and then, CTmax is recorded as the body temperature at which each ant has its hind legs paralyzed, disorganizing its locomotion.

**Figure 1.**
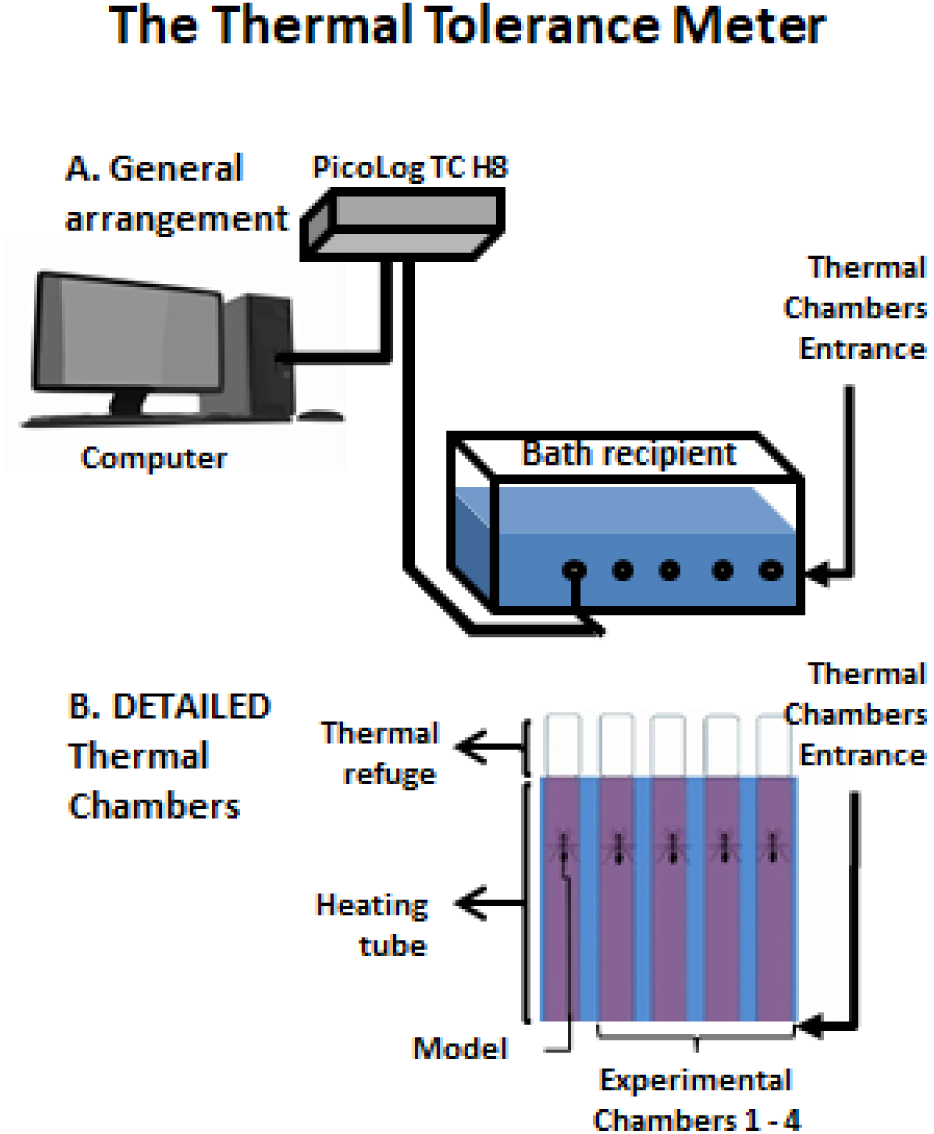
The Thermal Tolerance Meter. **A)** General view of all experimental arrangement **B)** In a detailed view is possible to see 5 chambers, 1 for the model and 4 experimental.

### 2.3. Experiments

#### 2.3.1. Effects of body size on VTM and CTmax

We measured the VTM and CTmax of 49 individuals with varying sizes (1.5-4 mm of head width). Size was measured with an analogic caliper (accuracy: 0.001m.). For this set of measurements, we used ST = 25°C and HR ~1°C/min.

#### 2.3.2. Combined effects of hydration level and relative humidity conditions on VTM and CTmax

The VTM and CTmax of individuals with hydration ranging 100-75 % were tested at environmental RH of 50% and 85%, always using individuals of 2.3mm in head width, ST = 25°C and HR ~1°C/min. The experiments were carried out in a controlled climatized room. The RH of the air and inside thermal chambers was similar, as checked using a hygrometer (HT-600 Instrutherm). Keeping probes measuring humidity within the thin thermal chambers was impossible, but even if some unnoticed variation in RH might happen during heating, it can be safely assumed that tubes at 50% remained always much drier than the ones at 85%.

#### 2.3.3. Combined effects of heating rate and start temperature on VTM and CTmax

Within our heating chamber, we heated ants using rates between 0.5°C/minute and 3.5°C/minute. Along these tests, we also varied ST between 23°C and 33°C. For this evaluation, we only used ants with 2.3 mm in head width, directly taken from the colonies.

### 2.4. Data analyses

We tested for all these effects, separately, fitting Linear Mixed Models (Bates et al., 2014) that related either the VTM or the CTmax with the described predictors. Thus, in each fitted model, either the VTM or CTmax was the response variable, and the corresponding predictors entered as fixed factors. The source colony for each ant entered as random factor to control for lack of independence in traits among sister ants.

When testing the effects of BS on the VTM and CTmax, we observed that average workers might present higher VTM than both, smaller and larger ones. Because of that, we obtained additional measurements on these same conditions and compared the fit of non-linear and linear relationships between ants’ size and VTM. We used the Akaike Information Criterion (AIC, Akaike, 1974) to choose the model with lowest values among three models: A first order one, or straight line, a second order one, describing a quadratic increase/decrease, and third order polynomial model, describing a parabolic trajectory.

When testing for the influence of the relative importance between HL and RH on VTM and CTmax, we used the Akaike’s information criterion to select between three competing models: (1) no effect, (2) interaction between HL and RH, (3) no interaction between them.

All analyses were performed in the R environment (R Development Core Team, 2018) using the package nlme (Pinheiro et al., 2017).

## 3 Results

### 3.1. Body size effect on VTM and CTmax

BS didn’t show linear correlations with VTM (N = 49, B: 0.13014, p: 0.6571) or CTmax (N = 49, B: 0.18545, p: 0.6214). However, a polynomial relationship of third order explained better the variation in VTM than a straight line, as indicated by a difference larger than 20 Akaike units (1st degree = 226,168, 2nd degree = 224,408 and 3rd degree = 205,112). That indicates that average-sized workers (with 2~2.6mm head width) consistently exhibited higher VTM, despite showing no differences in CTmax (Fig. 2A).

**Figure 2.**
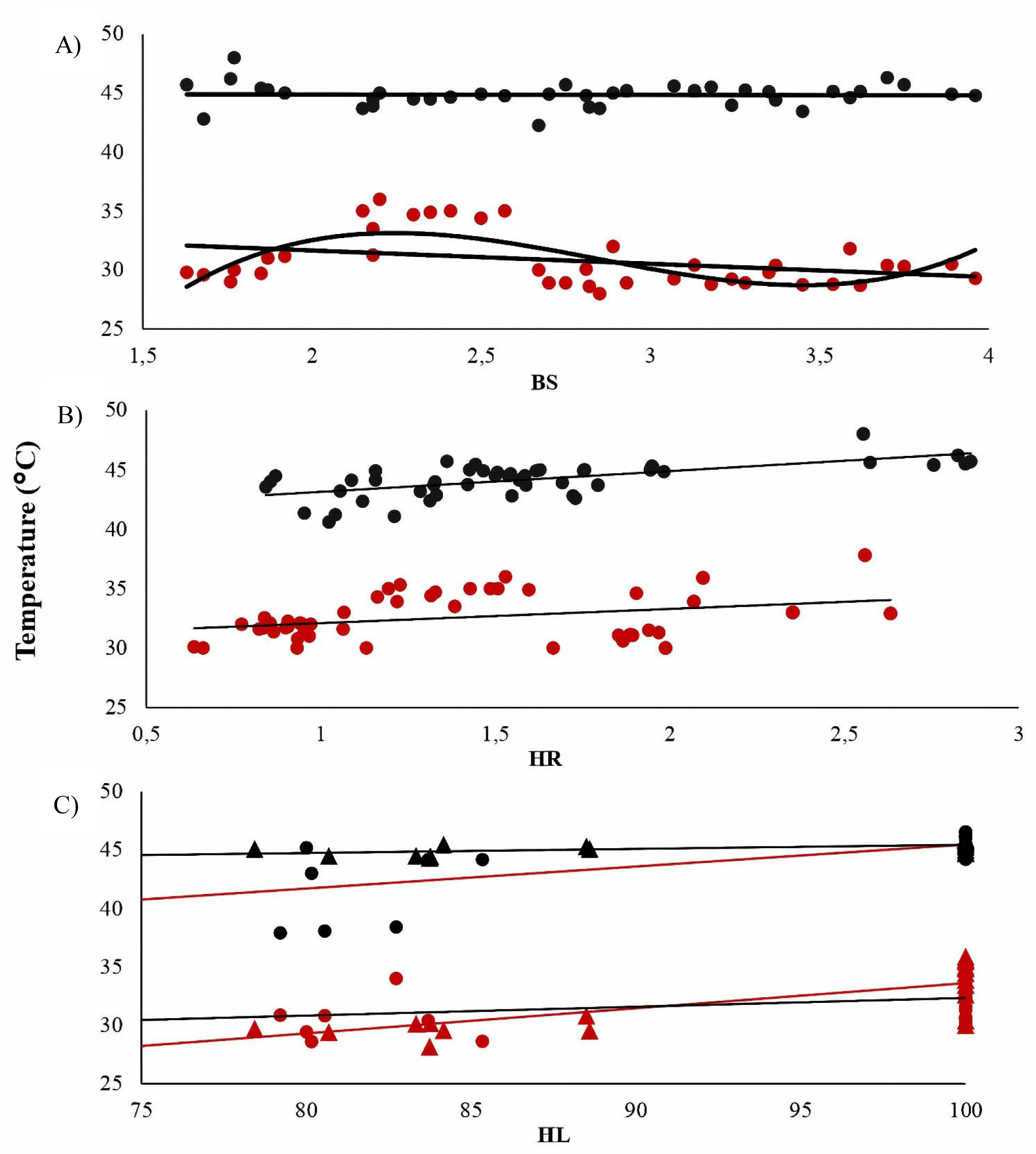
Relationships of ants’ VTM (in red) and CTmax (in black) with several factors. **(A)** While CTmax remained unaltered by body size (in mm of head width), the VTM of average-sized workers is higher than of their sisters; **(B)** Heating rates (in °C/minute) also increase both, the VTM and the CTmax linearly. Finally, (**C**) while body hydration level (in %) was important to increase both, the VTM and the CTmax, only the latter was altered by the relative humidity of the tubes. Black tendency lines and • represent the treatment at 50% relative humidity. Red tendency lines and ▴ represent the treatment at 85% relative humidity.

### 3.2. Effects of Heating rate and start temperature on VTM and CTmax

Raising the HR increased both, ants’ VTM (N = 57; B: 1.18781, p: 0.0073) and CTmax (N = 57; B: 1.77384, p: 0.0001), making VTMs range from 30°C to 37.8°C and CTmax from 40.6°C to 48°C (Fig. 2B). On the contrary, ST had no visible effects on either ants’ VTM (N = 45; B: 0.042469 p: 0.6653) or CTmax (N = 45; B: 0.00335, p: 0.9445).

### 3.3. Combined effects of hydration level and relative humidity on VTM and CTmax

Regarding the VTM, the full model without the interaction rendered the lowest AIC (7 units below the second one). According to it, only HL affected ants VTM, positively (N = 40, Nests= 5, B= 0.1571, SD= 0.02, t= 5.6, p: 0.000).

With respect to CTmax, the full model without interaction again rendered the lowest AIC (4 units below the second one). On this model, both hydration and RH had positive effects on CTmax, with relative umidity having stronger effects (Hydration: N = 40, Nests=5, B= 0.4129, SD= 0.0169, t= 2.4426, p: 0.020; Humidity: N = 40, Nests=5, B= 0.110853, SD= 0.0273145, t= 4.058415, p: 0.003) (Fig. 2C).

## 4 Discussion

Our most striking finding is that average-sized workers “dare” to get closer to their CTmax, compared to smaller or larger sister ants. Medium-sized individuals belong to a worker cast which spends more time outside the colony, foraging (Wilson, 1980). Thus, our results open the possibility that labor predisposition in ants, instead of being explained by body size alone (Wilson, 1980), might correspond with the development of specific tolerance thresholds (e.g. the VTM) that manage the probability of engaging in different tasks. Considering that neural dysfunctions precede physiological changes (Jørgensen et al., 2020), it is possible to speculate that behavioral and/or physiological aspects can be differentiated for small and large sister ants, along their ontogeny. Among hymenopterans, it is common that the separation of reproductive and working casts is supported by feeding with different substances (Dussutour and Simpson, 2009; Markin, 1970; Petralia and Vinson, 1978), but no similar mechanism has been yet discovered to explain for differences in thermal traits. In turn, having thermally daring workers might be rewarded with longer foraging times during hotter periods, and larger foraging areas, while the rest of the colony occupies cooler spaces. More thermotolerant species have been found to be more abundant among Mediterranean species (Cerdá et al., 2002). Yet, as a consequence of spending longer times foraging at higher temperatures, the lifespan of average workers might be shorter, compared to workers specialized in bringing water (smaller ants, Ribeiro and Navas, 2008) or defending the colony (the larger ones, Powell and Clark, 2004).

Our results raise methodological concerns on previous values of CTmax, for leaf-cutting ants and potentially other small arthropods. Often, previous studies have warmed ants using a heated ground surface while measuring the temperature right over it, with a probe. This approach generates strong thermal gradients between the ants’ body, the heating surface, and the measuring probe. In turn, our system provided homogeneous warming (Figure S5), and that prevented ants from creating a large thermal gradient when raising over their legs. Thus, our results do not support the idea that larger bodies render ants with higher thermal tolerance (Ribeiro et al., 2012), although longer legs might still protect larger ants from heating surfaces (Sommer and Wehner, 2012). Heating ants from below and measuring the temperature at the plate or right over it likely overestimates the CTmax of ants, or other arthropods. Supporting this idea, Ribeiro et al (2012) measured CTmax up to 53°C, while our maximum CTmax was 48°C.

While this difference does not invalidate their conclusions, the problematic approach may lead to largely underestimate the true thermal vulnerability of arthropods. When using large global physiological databases, like the GlobTherm (Bennett et al., 2019), such details are difficult to evaluate, as is the case for ant species. For example, *Pogonomyrmex desertorum* is considered one of the most thermophilic species known (CTmax = 53°C). In GlobTherm, the data entry is referred to a thesis by A.C. Marsh (1985). However, as others species with CTmax over 50°C, it was measured in such a problematic manner by Whitford and Ettershank (1975). In this way, our results exemplify the critical importance of homogenous warming during the determination of thermal tolerance in arthropods.

We also found that both, relative humidity and hydration level can raise the CTmax of leaf-cutting ants. Humidity also increased the CTmax of termites (Woon et al., 2018), while hydration has been observed to increase both, the CTmax and VTM of ectothermic vertebrates (e.g. Anderson & Andrade 2017; Guevara-Molina et al under review, Camacho et al under review). Therefore, hydration may be important for physiological thermal tolerance, across a variety of terrestrial ectotherms.

We know no previous studies that have compared internal and external hydric cues on maximum voluntary temperatures, but leaf-cutting ants seem to react only to internal cues (i.e. hydration level) rather than environmental cues signaling coming hydric stress. These experimental responses add up to previous findings of water stressed ants selecting leaves with higher water content (Bowers and Porter, 1981), and support the idea that behavior will buffer hydric stress in response to the internal hydration state. Future studies should test whether this explanation holds across different organisms, so we can better model their responses to climatic challenges that involving warming and droughts.

Nonetheless, ants adjusted their VTM and CTmax to environmental clues, such like heating rates. The VTM increased with heating rate, in parallel with their CTmax. Yet, total heating time did not, as shown by the lack of effect of the start temperature. Heating time might only become important at sublethal temperatures, and not at the long range studied herein (23°C ~ 30°C). At temperatures close to the VTM (32°C ~ 33°C), thermally enhanced processes, like production of free radicals (Luschack, 2011); or dehydration rates (Edney, 1977; Lighton and Bartholomew, 1988), may need time to accumulatively overwhelm homeostatic processes, or for triggering neurological signals of overheating. Adjustment of the VTM to heating rates also happens in at least some species of flies, lizards and amphibians (Terblanche et al. 2007; Camacho et al., 2018; Guevara-Molina et al under review, respectively). To continue understanding integrative responses of organisms to temperature rises, further studies could compare the power of different stressful conditions, such as time at sublethal temperatures, relative humidity, or oxygen availability. Basing these studies on serial measures of the VTM and CTmax would also allow to contextualize the behavioral response (VTM) on the effects that these factors have on the thermal limits.

Concluding, we observed how an ectotherm’s thermal tolerance responds to environmental and internal cues, in terms of both its behavior (VTM) and physiological limits (CTmax). Leaf-cutting ants became less “heat daring” when dehydrated or if heated more slowly, apparently reacting to an increased risk of overheating, signaled by a lowered CTmax, in both cases. Also, the non-linear effects of their body size on their VTM might have implications for their labor division. Many more studies will be necessary to correctly model how behavioral and physiological thermal tolerance integrates across species, but the urgency to make them is more pressing than ever, given the current climatic trends. With some adjustments, the device we developed should allow estimating the VTM and CTmax for other small ectotherms (e.g. bees, caterpillars, spiders, etc).

## List of abbreviations

BS: Body Size
CTmax: Critical Thermal Maxima
HL: Hydration Level
HR: Heating Rate
ST: Start Temperature
RH: Relative Humidity of the Air
VTM: Voluntary Thermal Maxima

## Acknowledgments

We thank FAPESP and CAPES to finance our study (FAPESP 2018/15664-5). We also thank Prof. Dr. Márcio Reis Custódio and Prof. Dr. Fernando Ribeiro Gomes for letting us conduct our experiments in their laboratory’s dependences.

## Competing Interest

The authors declare that the research was conducted in the absence of any commercial or financial relationships that could be construed as a potential competing interest.

## Author Contributions

Conceptualization: C. S. L., A. C., A. F. H.; Methodology: C. S. L., A. C.; Experiments: C. S. L., Resources: A. C., A. F. H.; Data curation: C. S. L., A. C.; Data analysis: A. C.; Writing: C. S. L., A. C.; Funding acquisition: C. S. L., A. C., A. F. H.

## Funding

This work was funded by Fundação de Amparo à Pesquisa do Estado de São Paulo (FAPESP grant 2018/15664-5 to C. S. L.). A. C. G. was funded by FAPESP (12/15754-8), CAPES (001) and Marie Curie Grant (897901) during the preparation of this study.

## Data availability

The raw data and supporting figures can be accessed here: 10.6084/m9.figshare.12762038

